# The diversity and functional capacity of microbes associated with coastal phototrophs

**DOI:** 10.1101/2022.01.05.475171

**Authors:** Khashiff Miranda, Brooke L. Weigel, Emily C. Fogarty, Iva A. Veseli, Anne E. Giblin, A. Murat Eren, Catherine A. Pfister

## Abstract

Coastal marine phototrophs exhibit some of the highest rates of primary productivity in the world. They have been found to host a diverse set of microbes, many of which may impact the biology of their phototroph hosts through metabolisms that are unique to microbial taxa. Here we characterized the metabolic functions of phototroph-associated microbial communities using metagenomes collected from 2 species of kelp (*Laminaria setchellii* and *Nereocystis luetkeana*) and 3 marine angiosperms (*Phyllospadix scouleri, P. serrulatus* and *Zostera marina*), including the rhizomes of two surfgrass species (*Phyllospadix* spp.) and the seagrass *Zostera marina*, and the sediments surrounding *P. scouleri* and *Z. marina*. Using metagenomic sequencing, we describe 72 metagenome assembled genomes (MAGs) that potentially benefit from being associated with macrophytes and may contribute to macrophyte fitness through their metabolic gene content. All host-associated metagenomes contained genes for the use of dissolved organic matter from hosts and vitamin (B_1_, B_2_, B_7_, B_12_) biosynthesis. Additionally, we found a range of nitrogen metabolism genes that transform dissolved inorganic nitrogen into forms that may be more available to the host. The rhizosphere of surfgrass and seagrass contained genes for anaerobic microbial metabolisms, including *nifH* genes associated with nitrogen fixation, despite residing in a well-mixed and oxygenated environment. The range of oxygen environments engineered by macrophytes likely explains the diversity of both oxidizing and reducing microbial metabolisms, and contributes to the functional capabilities of microbes and their influence on carbon and nitrogen cycling in nearshore ecosystems.

**Importance:** Kelps, seagrasses and surfgrasses are ecosystem engineers on rocky shorelines where they show remarkably high levels of primary production. Through analysis of their associated microbial communities, we found a variety of microbial metabolisms that may benefit the host, including nitrogen metabolisms and the production of B vitamins. In turn, these microbes have the genetic capability to assimilate the dissolved organic compounds released by their phototroph hosts. We describe a range of oxygen environments associated with surfgrass, including low-oxygen microhabitats in their rhizomes that host genes for nitrogen fixation. The tremendous productivity of coastal phototrophs is likely due in part to the activities of associated microbes and an increased understanding of these associations is needed.

## Introduction

We are experiencing a paradigm shift in biology with the recognition that many species exist as a consortium with microbes (1). These microbial associations are nearly ubiquitous, spanning a diversity of hosts across ecosystems. In coastal marine environments, phototrophic microbial hosts are diverse and range from marine angiosperms to large eukaryotic protists (macroalgae). Different macroalgal host species (2, 3) and different phototroph tissues (4, 5) host distinct microbial communities numbering in the millions per cm^2^ of host tissue (6), yet we still know little about the functional role the microbiome plays in host fitness or how the host influences the microbiome. The microbiome of phototroph species has been shown to have metabolisms that provide nitrogen to the host (7, 8). Bacteria also supply B vitamins (9) and affect development of their host (10). Further, the contributions that marine phototrophs make to host carbon and nitrogen cycling have largely ignored the role that microbes play. Even as we begin to describe their microbiome, we are discovering that environmental change affects these communities (11). For many of the foundational phototrophic species in the coastal ocean, our understanding of the diversity and role of their microbiome is nascent.

A unique aspect of host-associated microbes are the strong gradients in oxygen that they experience due to the biological activities of the host. The photosynthetic and respiratory activities of the host can generate a ‘phycosphere’ (12) where the host influences the physical environment experienced by microbes, sometimes over micron or mm scales. For example, the basal leaf meristem of the seagrass *Zostera* ranges from oxic to anoxic conditions over a scale of 300 microns when measured with oxygen microsensors (13). This range of oxygen concentrations likely selects for a diversity of microbial metabolisms in association with macrophytes.

Another factor important to the microbial metabolisms associated with coastal macrophytes is nutrient availability. In coastal systems, nitrogen can limit primary production and microbial associates that aid in accessing nitrogen might be selected. Microbial metabolisms that can increase the available dissolved inorganic nitrogen (DIN) for the host (14) include pathways that cleave carbon-nitrogen bonds to generate ammonium. This ammonification in biological systems can result from a diversity of hydrolases, including ureases and other enzymes that cleave C-N bonds (15). Further, microbes that fix atmospheric nitrogen have been discovered in an increasing number of taxa (8, 16), now recognized to include heterotrophic as well as phototrophic taxa (17–19). Nitrogen fixation was previously assumed to be restricted to nitrogen-poor environments, but has been quantified recently in systems thought to be nitrogen-rich (8, 20), an enigmatic finding given that nitrogen fixation is a costly metabolic process that consumes 16 ATPs per N_2_ fixed (21). Sediments where oxygen is low and nutrients can be depleted by macrophytes, such as the rhizosphere of seagrasses have provided evidence of nitrogen fixation (22–26). The recent discovery that nitrogen fixation takes place on particles in the coastal ocean where nitrate is relatively abundant (8, 20) suggests that *nifH* genes could be abundant in other nearshore systems.

Microbial metabolisms that synthesize compounds and vitamins needed by seaweeds and seagrasses may also underlie host-microbe exchanges. The active form of Vitamin B1 (thiamin) is essential for all organisms and is involved in carbohydrate and amino acid metabolisms. Vitamin B2 (riboflavin)-binding proteins are co-enzymes in various oxidases and are involved in photosynthesis and phototropism (27). Vitamin B7 (biotin) is a cofactor for acetyl coenzyme A (coA) which is essential for fatty acid synthesis. Vitamin B12 (cobalamin) is required as a coenzyme in the mitochondria for many algae, yet they depend upon prokaryotes to produce it (9, 28). Thus, marine macrophytes may be auxotrophic for key vitamins, and their production by host-associated bacteria may be another basis for phototroph-microbiome interactions in nature.

Hosts might reciprocally benefit microbes, especially if heterotrophic microbes benefit from the dissolved organic carbon that is released by their hosts. Of the carbon that is fixed, kelp have been demonstrated to release 15-16% of it as dissolved organic matter (29, 30), and seagrasses too provide a constant source of dissolved organic carbon (31, 32), likely stimulating heterotrophic bacterial processes (33). These rates of organic carbon release, often involving highly labile organic carbon compounds (34), could provide the basis for reciprocal benefits between microbes and their associated hosts.

Here, we analyzed microbial metagenomes collected from 5 different coastal phototrophs to determine if there is functional genomic evidence of microbial metabolisms that could reciprocally benefit host and microbes. We analyzed the surface microbiome on the blade of two kelp species (*Laminaria setchellii* and *Nereocystis luetkeana*) and the surfgrass *Phyllospadix scouleri*, the rhizomes of *P. scouleri, P. serrulatus*, and the seagrass species, *Zostera marina*, and the sediment surrounding the rhizomes of *P. scouleri* and *Z. marina*. We quantified the variable oxygen environment in the rhizomes of *Phyllospadix spp*. to determine if they allow for aerobic as well as anaerobic metabolisms. We analyzed the microbial taxa present and examined their gene content to estimate their functional and metabolic capacities. We hypothesized that microbial partners: 1) enhance host access to dissolved inorganic nitrogen through nitrogen recycling, ammonification and nitrogen fixation, 2) provision vitamins B_1_, B_2_, B_7_, B_12_, and 3) use a diversity of abundant dissolved organic carbon exudates from the host. We tested whether microbial taxonomy and function differed across hosts and host tissue types, and whether anaerobic metabolisms were present in low-O_2_ environments (e.g., rhizomes and sediment). Through this study, we find that the range of oxygen environments engineered by host phototrophs likely explains the diversity of both oxidizing and reducing microbial metabolisms, and contributes to the functional capabilities of microbes and their influence on carbon and nitrogen cycling in nearshore ecosystems.

## Methods

### Sampling and DNA Extraction

We collected metagenome samples from the surfaces of 5 different phototroph species (Table S1). The surface of *Phyllospadix scouleri* blades, *Laminaria setchellii* fronds and the inner bulbs of *Nereocystis luetkeana* were swabbed with a sterile swab and brushed with an interdental brush (GUM Proxabrush Go-Betweens). We preserved sections of the rhizomes of *Phyllospadix scouleri, P. serrulatus* and *Zostera marina*. Sediment surrounding *P. scouleri* and *Z. marina* was also collected. All samples were collected from Tatoosh Island, WA, USA (48.393679, - 124.734617) on 16-17 Jul 2019, except for *Z. marina* samples which were sampled from West Falmouth Bay, MA, USA (41.60708333, −70.64527778) on 19 Sept 2019. We included samples from the rhizosphere of *Z. marina* from the Atlantic Ocean as a known positive control for nitrogen fixation (22, 23). Swabs, tissue and sediment were immediately frozen at 20° C and shipped to storage at −80° C. DNA from these collections was extracted with a Qiagen PowerSoil Kit and multiple samples were pooled for each metagenome sample to increase DNA quantity and possible discovery: *P. scouleri* blade, rhizome and sediment (3 pooled individuals each), *P. serrulatus* rhizome (3 individuals), *L. setchellii* blade (3 individuals), *N. luetkeana* interior bulb (4 individuals), *Z. Marina* rhizomes and sediment (2 individuals).

### Shotgun metagenomic sequencing, assembly, and read recruitment

The above 8 samples were run over 2 lanes on a HiSeq 2500 (2×150) with TruSeq DNA library preps at Argonne National Laboratory. For each sample, resulting DNA sequences were first quality filtered (35)(Minoche et al. 2011), then assembled with IDBA-UD v1.1.3 (36) (Peng et al. 2012) with a minimum scaffold length of 1 kbp. Metagenomic short reads from each sample were then recruited back to their corresponding assembled contigs using Bowtie2 (37). Samtools (38) was used to generate sorted and indexed BAM files. Anvi’o v7.0 (39) was used as the command line environment for all downstream analyses. ‘anvi-gen-contigs-database’ was used to generate anvi’o contigs databases, during which Prodigal v2.6.3 (40) identified open reading frames, and ‘anvi-run-hmms’ was used to identify genes matching to archaeal and bacterial single-copy core gene collections using HMMER (41).

### Reconstructing metagenome-assembled genomes (MAGs)

To reconstruct genomes from the assembled metagenomes, we used a combination of automatic binning via CONCOCT v1.1.0 (42), followed by a manual curation of each MAG as outlined by Shaiber et al. 2020 (43). Genome taxonomy was determined using GTDB v.1.3.0 (44), and ‘anvi-run-scg-taxonomy’. We also inferred gene-level taxonomy using Centrifuge v1.0.4 (45) to aid manual curation.

### Phylogenomic analysis of MAGs

To perform a phylogenomic analysis of our MAGs, we recovered amino acid sequences for bacterial single-copy core genes (SCGs) from each genome (except the only archaeal genome in our collection) using the program ‘anvi-get-sequences-for-hmm-hits’ with the parameter ‘--hmm-source ‘Bacteria_71’ on the ribosomal gene set ‘Ribosomal L1-L6’ and the flag ‘--concatenate’, which independently aligned each SCG independently using Muscle v3.8.1 (46) before concatenating them into a final superalignment. We then refined the alignment using trimAl v1.4.rev15 (47) to remove any position in the alignment if more than 50% of the residues were gap characters. A maximum-likelihood phylogeny was inferred using IQTree (48) with 1,000 bootstrap replicates, and a LG+R6 model best fit our data using ModelFinder (49).

### Functional analysis of microbial communities

To address the metabolic capabilities of host-associated microbes, we annotated genes in each anvi’o contigs database with 3 different databases using ‘anvi-run-kegg-kofams’, ‘anvi-run-ncbi-cogs’, and ‘anvi-run-pfams’, which used the databases of Kyoto Encyclopedia of Genes and Genomes (KEGG) (50), NCBI’s Clusters of Orthologous Genes (COGs) (51) and EBI’s Pfam database (52) respectively. We used these annotated genes to test for 1) nitrogen cycling metabolisms, especially those within the nitrogen-fixation pathway, 2) hydrolases, including ureases, as well as ammonia-lyases, to cleave the C-N bonds in amino acids and make ammonium available to the host, 3) vitamin production, namely vitamins B_1_, B_2_, B_7_ and B_12_ and 4) a set of dissolved organic matter (DOM) transporter genes identified by Poretsky et al. (34) that indicate the ability of the microbial community to assimilate DOM exudates from kelps and surfgrasses. The list of genes used is indicated in Table S4. We additionally developed and used a graph-based algorithm on KEGG definitions for vitamins B_1_, B_2_, B_7_ and B_12_ to detect the presence of these biosynthetic pathways (Supplementary Code 1). To expand our functional analysis of kelp blade genes, we included 32 MAGs from the surface of *N. luetkeana* blades that were collected from the same location at the same time using similar methods as those described above (53).

### Phylogenetic analysis of nifH genes

To search for *nifH* amino acid sequences in our environmental samples, we identified 9 MAGs which contained *nifH* genes using the KEGG identifier K02588 with e-value < 1e-20. We aligned the AA sequences for these genes against 89 well-characterized reference *nifH* AA sequences (Table S6) using Muscle v3.8.1 (46) and refined the alignment using trimAl (gap-threshold: 0.5) and ‘anvi-script-reformat-fasta’ (max-percentage-gap: 50%). A maximum-likelihood phylogeny was inferred using IQTree (48) with 1,000 bootstrap replicates, and a LG+R5 model best fit our data using ModelFinder (49). *nifH* genes from the *Zostera* samples served as positive controls to detect nitrogen fixation genes in other samples. Figures 2, 3, 4 were generated using iTol v5 (54), R v4.0.3 and FigTree respectively. We additionally took tissue samples from *P. scouleri* rhizome (n = 16), basal meristematic region just distal to the sheath (n = 12) and blade 35 cm above the rhizome (n = 12) to quantify stable isotopes of δ15N and δ13C to look for signatures of nitrogen fixation (methods described in Appendix 1).

**Figure 1.**
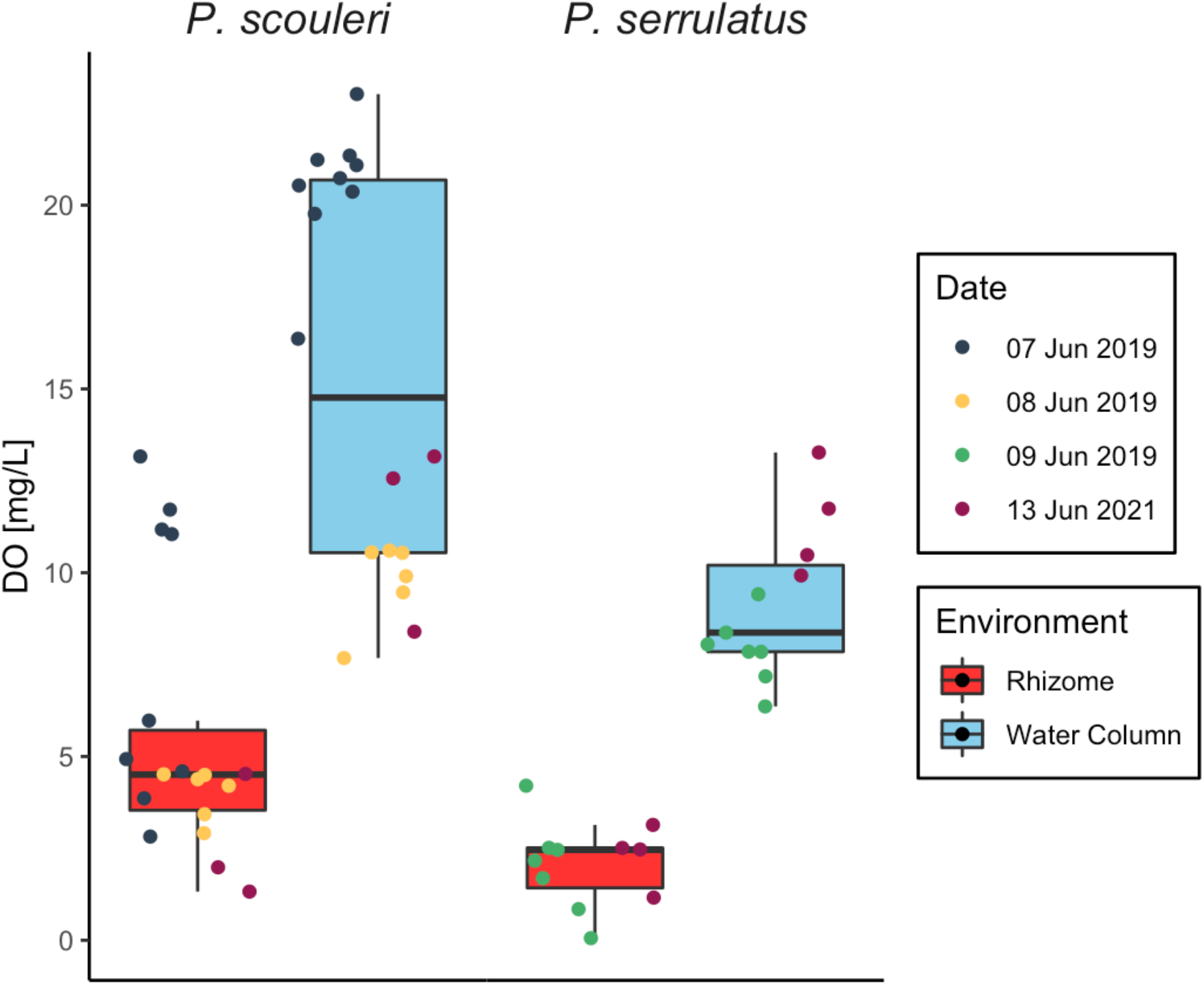
Boxplot comparing the dissolved oxygen concentrations of water column (blue) and the sediment-rhizome environment (red) of *P. scouleri* (pairwise t-test: p<0.001) and *P. serrulatus* (pairwise t-test: p<0.001). Sampling dates are represented by different colors.

**Figure 2.**
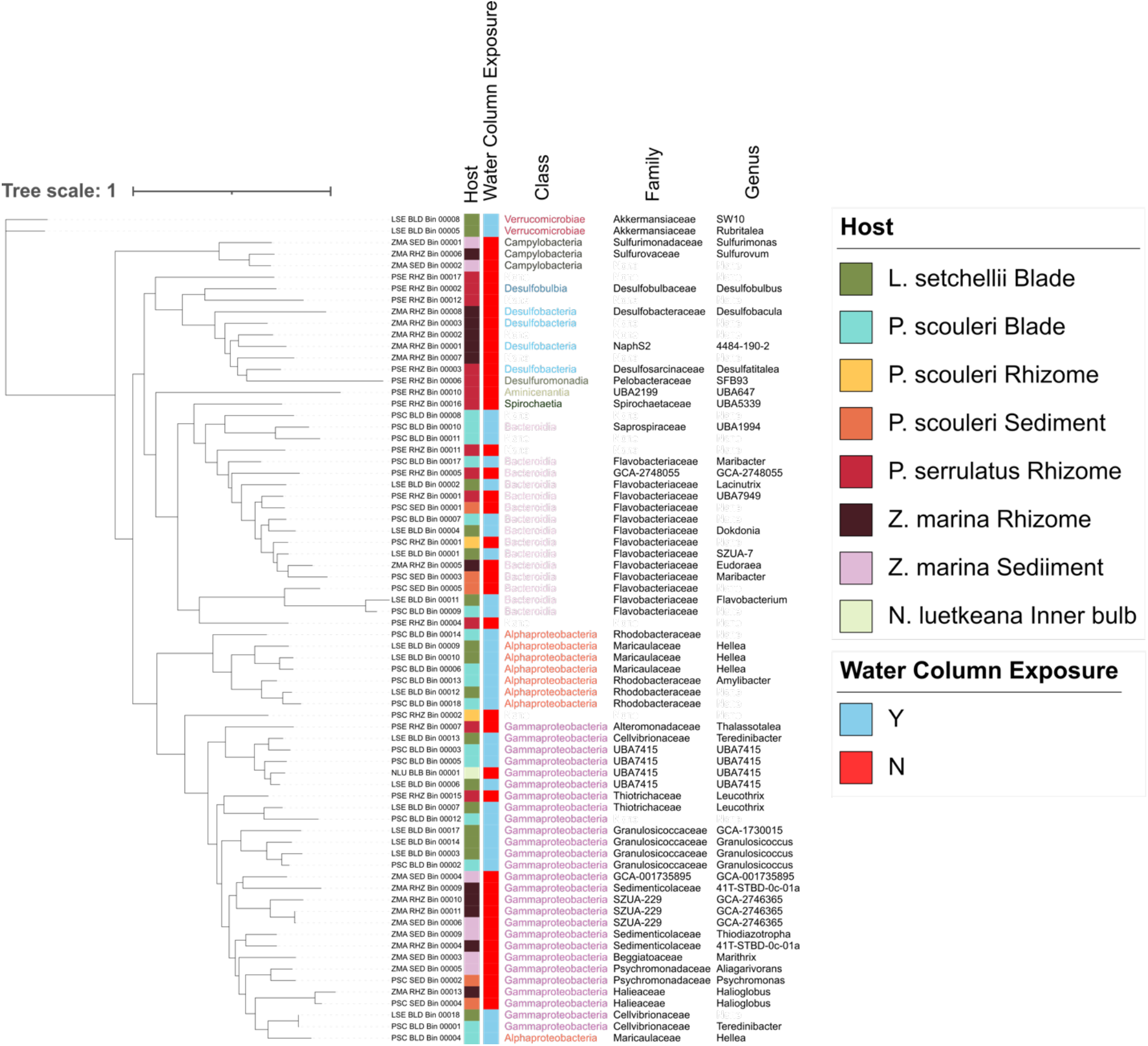
A phylogenomic tree of 6 concatenated bacterial single-copy core ribosomal genes from 71 bacterial MAGS across 8 samples, showing the results from 33 high quality MAGs and 38 lower quality ones. One MAG, PSC_RHZ_Bin_00003, from the rhizome of *P. scouleri,* was identified as an archaeal genome and was thus omitted from this tree. Gaps in class, family and genus indicate the level to which taxonomic classification was resolved in each MAG. All blade tissues have ‘water column exposure’, while rhizome and sediment samples do not.

**Figure 3.**
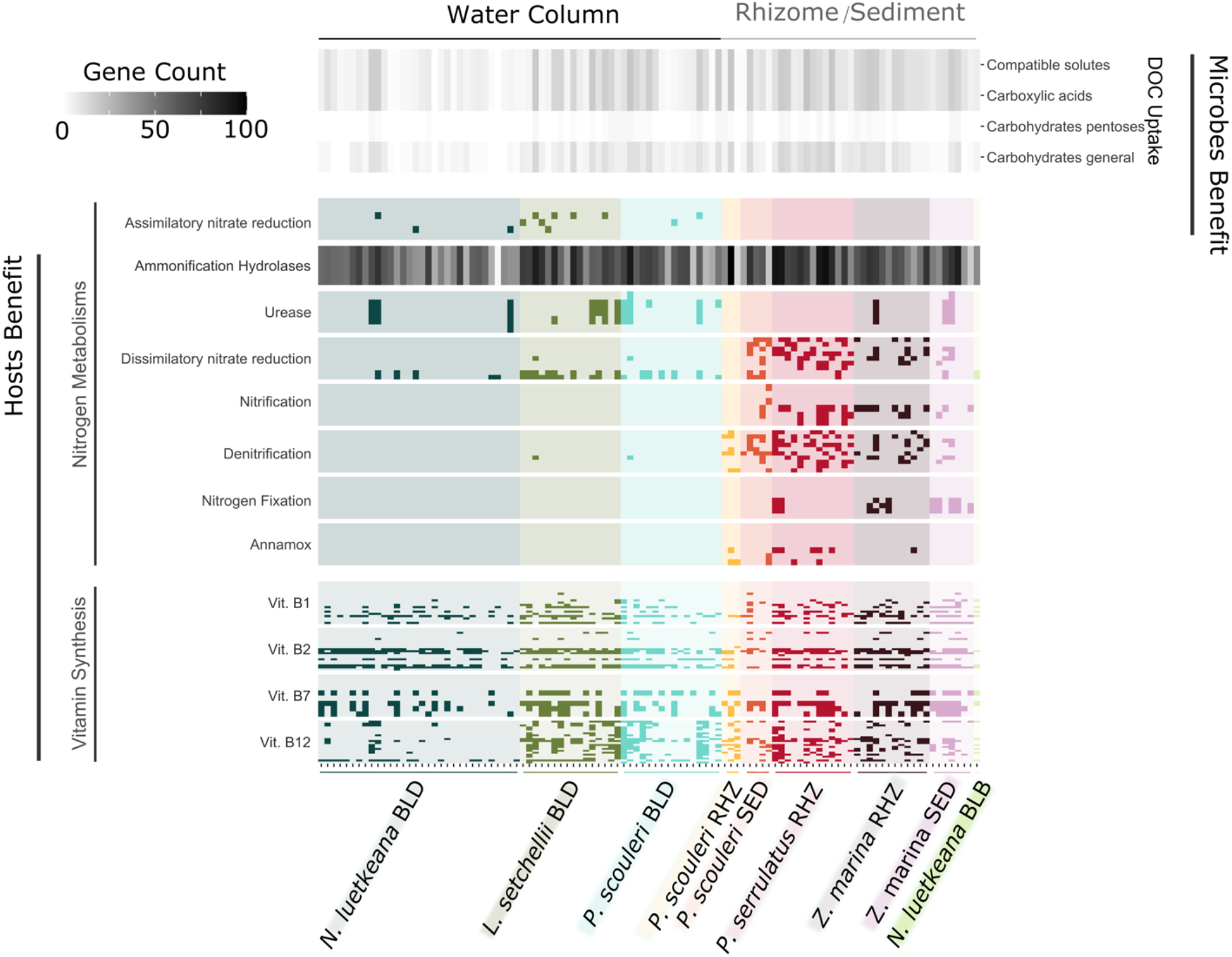
Microbial Metabolisms in the MAGs reported in Fig. 2 and Table S3 across all hosts and grouped as those that might benefit the host (“hosts benefit”) and microbial metabolisms that might utilize host provisioned metabolites (“microbes benefit”). Each tick along the x-axis corresponds to a MAG. *N. luetkeana* blade MAGs are from Weigel et al. (in review). The metabolisms for **DOC Uptake** that benefit microbes are shown as a heatmap of the count of the number of genes that can metabolize *compatible Solutes, carboxylic Acids, carbohydrate Pentoses* and *General carbohydrates*. Microbial metabolisms that benefit the host are **Ammonification Hydrolases**, where the heatmap provides a count of the hydrolases acting on C-N bonds other than peptide bonds, **Nitrogen Metabolisms** and **Vitamin** Synthesis, both shown as the presence or absence of a gene in a pathway. The genes used in this are in Table S4.

**Figure 4.**
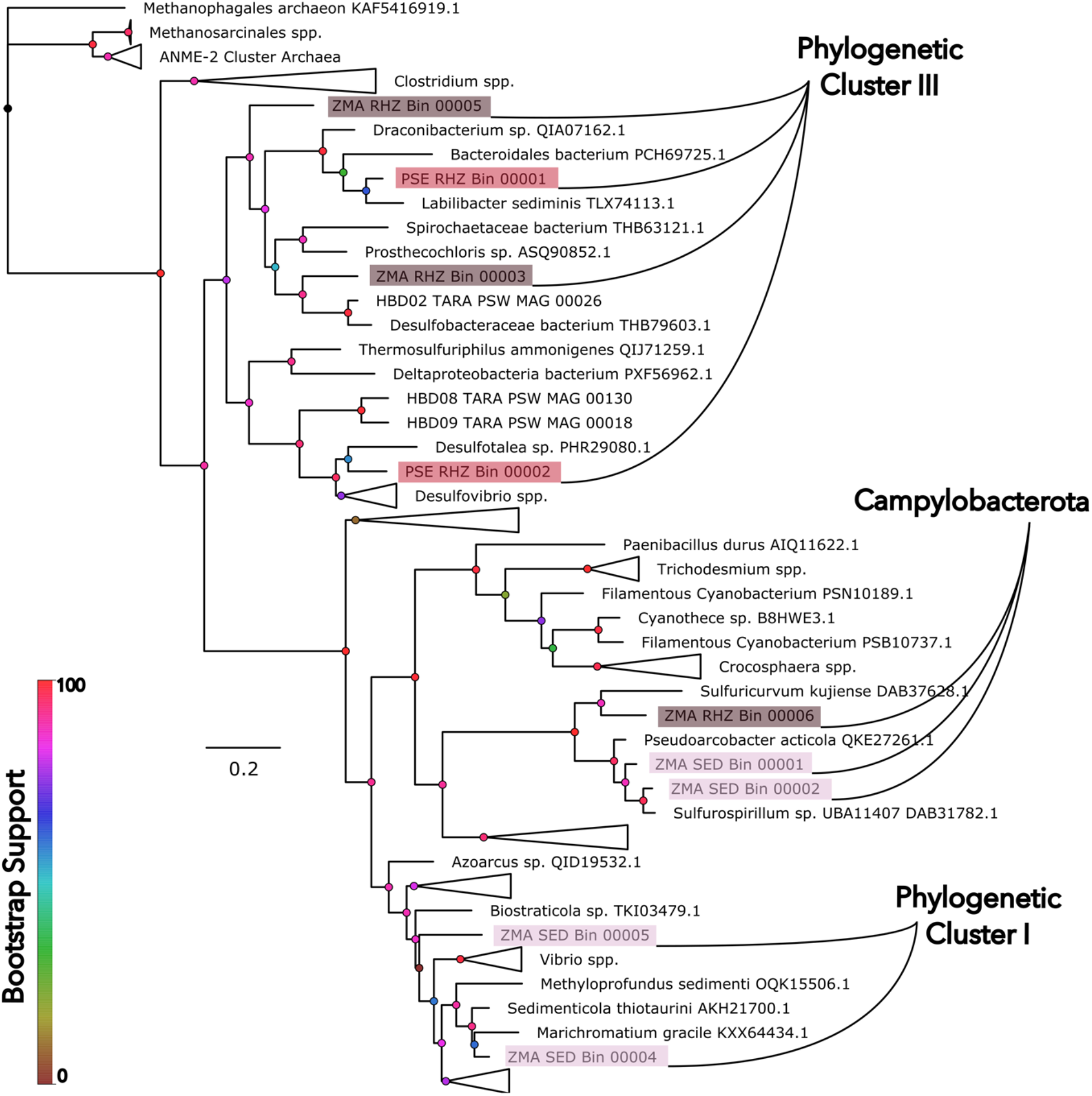
A phylogenomic tree of *nifH* genes found on the rhizomes of *P. serrulatus* (PSE, n = 3) and the rhizomes and surrounding sediment of *Z. marina* (ZMA, n = 2 and 5, respectively). Some *nifH* genes group into Cluster I, including a sulfur oxidizing taxon on the rhizome of *Z. marina*, and other taxa in *Campylobacterota,* including *Sulfurovum*. Cluster III contains taxa associated with rhizomes including rhizomes including *Desulfobulbus mediterraneus* on *P. serrulatus* and a Desulfobacterales associated with *Z. marina* rhizomes.

### Quantifying the Oxygen Environment

We quantified the oxygen concentrations in proximity to *Phyllospadix spp.* rhizomes by comparing dissolved oxygen (DO) concentrations in the surrounding seawater and in the sediment around the rhizome. We used a Pyro Science Robust Oxygen Probe (OXROB10, Firesting™, Pyroscience), and repeated measurements around 0900h across 4 days (7-9 June 2019, 13 June 2021) within *P. scouleri* (n = 18) and *P. serrulatus* (n = 11) rhizomes. Each reading first measured the surrounding seawater after which we gently pushed the tip of the oxygen probe into the sediment and rhizome mass to a depth of 1-3 mm, the typical thickness (*pers. observation*). We let the probe equilibrate and took a reading at 150 sec. This allowed the rhizome oxygen environment to equilibrate after we disturbed the intact rhizome. We compared surrounding water and within-rhizome oxygen using paired t-tests in R.

### Data Availability

In addition to the code available on GitHub (___), the final MAG database files generated in anvi’o are available on the FigShare repository: (_____). Metagenomic sequence data are available at the NCBI’s Sequence Read Archive under accession no. (submission in progress).

## Results

### Surfgrass rhizomes have lower oxygen concentrations than surrounding seawater

The oxygen environment in the rhizomes differed significantly from that of the surrounding seawater (Fig. 1). Rhizomes maintained a lower dissolved oxygen (DO) concentration than the surrounding seawater for both *P. scouleri* (n=18, pairwise t-test: *p*<0.001) and *P. serrulatus* (n=11, pairwise t-test: *p*<0.001). *P. serrulatus* maintained a slightly lower DO concentration in the rhizome at 2.11 mg l^-1^, compared to 5.61 mg l^-1^ for *P. scouleri*. However, the nature of sampling likely introduced more oxygenated water from the surrounding water column to the rhizome-sediment microenvironment, suggesting that the actual DO concentration within the sediment is lower than the value reported.

### Diversity of MAGs assembled across hosts

Following filtering, we obtained an average of 41 million reads per sample (range 6.48 to 67.73 million), with 79.8% of raw reads retained on average. When these reads were assembled into contigs of at least 1000 nucleotides, a mean of 42,026 contigs and a mean of 110,054 genes were present across samples (Table S1).

Across 8 metagenomes we manually binned 33 high quality MAGs, defined as having a completion score >90% and contamination (or redundancy) < 10% (Table 2). We also identified 39 lower quality MAGs that had completion scores between 38 and 93% and redundancy scores between 0 and 21% (Table S3). All MAGs were bacterial except for a single archaeal MAG on the rhizome of *P. scouleri*. The bacterial MAGs spanned 7 phyla, including *Proteobacteria* (n=34), *Bacteroidota* (n=19), *Verrumicrobia* (n=2), *Campylobacterota* (n=3), *Desulfobacterota* (n=5), and a single MAG in each of *Desulfomonadota, Acidobacteriota*, and *Spirochaetota*. The Archaea belonged to the phylum *Chrenarchaeota.* There were 46 MAGs resolved to the species level, with 8 to the genus level, 9 to family, 2 to order, and 2 to class level. Five MAGs were resolved only as Bacteria (Table S2).

**Table 1.** Summary of the features of 8 metagenomes. More information is in Table S1 and the taxonomy based on single copy genes is in Table S2.

| *Phyllospadix*<br>*scouleri* |  |  | *Phyllospadix*<br>*serrulatus* | *Laminaria*<br>*setchellii* | *Nereocystis*<br>*luetkeana* | *Zostera*<br>*marina* |  |
| --- | --- | --- | --- | --- | --- | --- | --- |
| Sediment | Rhizome | Blade | Rhizome | Blade | Inner bulb | Sediment | Rhizome |
| # quality reads (in millions) |  |  |  |  |  |  |  |
| 43.68 | 67.73 | 38.41 | 37.99 | 48.58 | 6.48 | 19.37 | 65.76 |
| Bacteria |  |  |  |  |  |  |  |
| 63.7% | 58.3% | 63.6% | 63.0% | 63.9% | 62.2% | 33.6% | 60.6% |
| Archaea |  |  |  |  |  |  |  |
| 34.2% | 38.3% | 33.1% | 33.3% | 32.7% | 35.9% | 62.1% | 34.2% |

**Table 2.** Metagenome assembled genomes across all samples and their representation across phyla. More detailed information on the MAGs can be found in Table S3.

| *Phyllospadix*<br>*scouleri* |  |  | *Phyllospadix*<br>*serrulatus* | *Laminaria*<br>*setchellii* | *Nereocystis*<br>*luetkeana* | *Zostera*<br>*marina* |  |
| --- | --- | --- | --- | --- | --- | --- | --- |
| Sediment | Rhizome | Blade | Rhizome | Blade | Inner bulb | Sediment | Rhizome |
| High Quality MAGs |  |  |  |  |  |  |  |
| 3 | 1 | 7 | 6 | 9 | 1 | 2 | 5 |
| Other MAGs |  |  |  |  |  |  |  |
| 2 | 2 | 8 | 7 | 7 | 0 | 5 | 7 |
| Proteobacteria |  |  |  |  |  |  |  |
| 2 | - | 9 | 2 | 10 | 1 | 5 | 5 |
| Bacteroidota |  |  |  |  |  |  |  |
| 3 | 2 | 5 | 4 | 4 | - | - | 1 |
| Verrucomicrobia |  |  |  |  |  |  |  |
| - | - | - | - | 2 | - | - | - |
| Campylobacterota |  |  |  |  |  |  |  |
| - | - | - | - | - | - | 2 | 1 |
| Desulfobacterota |  |  |  |  |  |  |  |
| - | - | - | 2 | - | - | - | 3 |
| Desulfuromonadota |  |  |  |  |  |  |  |
| - | - | - | 1 | - | - | - | - |
| Acidobacteriota |  |  |  |  |  |  |  |
| - | - | - | 1 | - | - | - | - |
| Spirochaetota |  |  |  |  |  |  |  |
| - | - | - | 1 | - | - | - | - |
| No ID |  |  |  |  |  |  |  |
| - | - | 1 | 2 | - | - | - | 2 |
| Crenarchaeota |  |  |  |  |  |  |  |
| - | 1 | - | - | - | - | - | - |

The 72 MAGs belong to diverse microbial phyla, which were distributed across the 5 host species and tissue types (Fig. 2). In some cases, bacterial taxa from kelp blade tissues were most closely related to bacteria collected from the rhizome or sediment of a seagrass, suggesting that closely related bacterial taxa can associate with diverse hosts. Known anaerobic sulfur cyclers like *Desulfobulbia, Desulfobacteria, Desulfuromonadia* and *Campylobacteria* (*Sulfurovum sp000296775* and *Sulfurimonas autotrophica*) were exclusively found in the low oxygen rhizome and sediment samples of *Zostera marina* and *Phyllospadix spp*. Conversely, *Alphaproteobacteria*, were exclusively found on surfaces exposed to the water column. *Gammaproteobacteria* was the only class found across the range of tissue types (6 out of 8 host environments). We did not include the only well-resolved archaeal taxon found in our samples, *Crenarchaea* (*P. scouleri* rhizome), as our analysis compared single-copy core genes specific to bacterial phyla.

### Host-associated microbial genomes contain pathways to synthesize vitamins, recycle nitrogen, and use host-generated carbon

We found evidence for a number of metabolic pathways that are likely important for exchanges between host phototrophs and their microbial partners (Fig. 3). Microbes on hosts had genes for diverse carbohydrate and carboxylic acid assimilation via cell membrane transport proteins. Host-associated microbes also had genes for a diversity of nitrogen metabolisms, including ureases and hydrolases that could regenerate ammonium. Nitrogen metabolisms were most diverse in rhizome and sediment samples where we identified both oxidizing (nitrification) and reducing (nitrate reduction, nitrogen fixation, denitrification) metabolisms, as well as metabolisms that both oxidize and reduce (annamox).

Every sample had at least one gene from B-vitamins biosynthesis pathways. Using a simple-path based algorithm on KEGG definitions (Supplementary Code 1), we determined that all microbial communities had the metabolic pathways to synthesize vitamins B_1_ (with the exception of the *P. scouleri* rhizome), B_2_ and B_7_ (except inside the bulb of *N. luetkeana*). The Vitamin B_12_ anaerobic biosynthesis pathway, however, was only present in MAGs found on the blades of *L. setchellii* (2) and *P. scouleri* (3) and the rhizomes of *P. serrulatus* (2) and *Z. marina* (1). Additionally, all three MAGs on the blade of *P. scouleri* that had this anaerobic pathway had the genes necessary to synthesize Vitamin B_12_ aerobically as well.

### *Novel detection of* nifH *genes in surfgrass*

We identified the nitrogenase gene (*nifH*) in 9 MAGs with e-value support < 1.3e-120 (KEGG) and < 1.1e-135 (COG). These 9 MAGs were assembled from *P. serrulatus* rhizomes (n = 2) and *Z. marina* rhizomes (n = 3) and the surrounding sediment (n = 4). Of these 9 MAGs, 5 were resolved to the genus level, while others were resolved to the order and family level, including *Campylobacterales, Desulfobacterales* and 2 *Flavobacteriaceae* (Fig. 4, Table S5). *nifH* genes identified in the rhizomes of *P. serrulatus* and *Z. marina* belonging to the class *Desulfobacteria* and family *Flavobacteriaceae*, clustered within Cluster III: anaerobic nitrogen-fixers that are often coupled with sulfate-reduction metabolisms. Samples from *Z. marina* sediment and rhizome also contained 3 *nifH* genes in *Campylobacterial* MAGs that clustered together in a sister clade to the aerobic nitrogen-fixers of Cluster I. The COG gene identified as *nifH* (COG1348) also includes the homologous protochlorophyllides, which are involved in photosynthetic pigment synthesis but have high sequence similarity to the *nifH* gene (21, 55). Instead, we used the KEGG gene (K02588) that does not detect these homologs. When we inspected genes on the same contig with *nifH*, we found a number of genes related to nitrogen fixation (Table S5), including *nifD* (COG 2710) in 7 of the 9 contigs, nitrogen regulatory protein PII (COG 347), *nifB* (COG 535), and multiple iron containing proteins including ferrodoxin and Fe-Mo cluster-binding proteins (Table S5).

## Discussion

### Phototroph tissues and sediment host distinct microbial taxa and functions

The phototroph species we sampled in this study are foundational in coastal ecosystems (56–59), yet a description of the diversity and function of their microbiomes have been lacking. All MAGs were bacterial, except for a single archaeal MAG (*Crenarchaeota*) in the rhizome of *Phyllospadix scouleri*, which was identified as *Nitrosopumulis*, a genus associated with nitrification (Table S3). Together, these 5 phototrophs hosted bacteria from 9 phyla. The only low diversity sample was the interior of the bulb of *Nereocystis*, where we assembled only a single MAG (*UBA7415 sp002470515*) suggesting that this environment of high carbon monoxide and nitrogen gas (60) may inhibit microbial activity or pose a highly selective environment. Blades of kelp and surfgrass, in contrast, were a locus of microbial diversity and function, a finding that is similar to many recent studies of macroalgal and seagrass microbiomes reporting high microbial diversity (2, 4, 5, 61-63). The functional attributes of microbial taxa associated with marine macrophytes include pathogen resistance (64), the ability to provision the host with B vitamins (9), and enhanced host algal fitness (65), perhaps through some of the nitrogen metabolisms we documented here (14, 66).

### Host-microbe interactions in a dynamic oxygen microenvironment

Grouping MAGs by microbial metabolisms (Fig. 3) showed key functional differences among phototroph hosts. Blade tissues that interacted directly with the water column were associated with microbial nitrogen metabolisms that were mostly oxidizing. The abundance of dissolved organic carbon from phototroph hosts (29–31, 59) might select for heterotrophic metabolisms. Indeed, we found an abundance of genes for dissolved organic matter assimilation and transport in all metagenomes, suggesting that hosts may stimulate heterotrophy in their associated microbial community similar to findings by Poretsky et al. (34). Improved characterization of the components of dissolved organic matter and the genomes of hosts will allow us to better assess complementarity in resource supply by hosts and resource use by microbes.

The host tissue types in this study differed in surface oxygen concentrations. Blade tissue interacts with the water column and is likely more oxygenated than rhizome tissue or sediments, though a previous study suggests there can also be a 60% reduction in oxygen along the immediate surface of kelp blades (67), and along the mucus layer where some kelp-associated bacteria reside (6). Over two-thirds of the bacterial taxa on blades of *N. luetkeana* belonged to families associated with obligately aerobic metabolisms, demonstrating the role of oxygen in shaping phototroph-associated microbial communities (68). The sediment surrounding the rhizomes of *Phyllospadix spp*. contained low oxygen microenvironments (Fig. 1) likely maintained by macroinvertebrate respiration (69)(Moulton and Hacker 2011), similar to the biological processes in the anaerobic sediment surrounding *Zostera* (13). Low rhizosphere oxygen concentrations likely structured the taxonomic composition of *Z. marina* to include anaerobic taxa such as *Campylobacteria*, *Desulfatitalea* and *Desulfobulbus*. The presence of anaerobes like *Desulfuromonadia, Desulfobaceria, Spirochaeta* and *Aminicenantia* in *P. serrulatus* rhizomes suggests sulfate reduction also occurs, possibly coupled to dissolved organic carbon use as an energy source (e.g. (70) Howarth & Hobbie 1982). Additionally, *Campylobacteria* and the genus *Thiodiazotropha* were associated with *Z. marina* and may remove detrimental sulfide accumulation through sulfur oxidation (71, 72).

Nitrogen metabolisms that were both oxidizing and reducing were found in MAGs associated with rhizomes of both *Z. marina* and *Phyllospadix* (Fig. 3), suggesting the potential for temporal niches when, for example, ammonium oxidation to nitrate occurs during high-O_2_ daylight periods, followed by nitrate reduction or nitrogen fixation during O_2_-depleted nighttime hours. Additionally, all MAGs in this study contained hydrolases that cleave carbon-nitrogen bonds to produce ammonium (14), recycling nitrogen compounds for host uptake. Oxidizing and reducing metabolisms are likely separated only by microns in the hosts studied here.

We detected biosynthetic pathways for vitamins B_1_, B_2_, B_7_ and B_12_ that are required by the auxotrophic phototroph hosts in this study (9, 10, 73). We found that only the blades of *P. scouleri* harbored MAGs with both anaerobic and aerobic biosynthetic pathways for Vitamin B_12_, suggesting that the variable oxygen environment driven by host-metabolism creates diverse metabolic niches for associated microbes. Strong gradients in oxygen and metabolically diverse microbial metabolisms are present in a diversity of animal hosts such as corals and sponges as a result of host metabolism (74-76). Fluctuating oxygen microenvironments might also promote cross-feeding, where microbial taxa produce a metabolite that can be consumed by other taxa. Cross-feeding is potentially important for nitrogen (77) and carbon metabolisms (78, 79) in microbial communities.

### Characteristics of previously undescribed nitrogen fixation in surfgrass

Building on recent studies that illustrate the association of nitrogen fixing microbes with a diversity of macroalgae (80) and seagrasses (22, 23, 81, 82), we found a previously undescribed diversity of nitrogenase genes associated with the surfgrass *Phyllospadix*. We detected *nifH* genes in *P. serrulatus* rhizomes that resolved into the Cluster I group of nifH genes, which are characterized by aerobic nitrogen fixers. *P. serrulatus*, in comparison to *P. scouleri*, is found higher up in the intertidal zone and often in sheltered tidepools that tend to undergo dramatic daily fluctuations in oxygen, possibly allowing for a temporal low-O_2_ niche during the night (83). Conversely, we did not detect nitrogenase genes in the microbiome of *P. scouleri*, which inhabits more wave-exposed and thus better oxygenated environments (Fig. 1). However, stable isotope analyses across *P. scouleri* samples show a lower nitrogen isotopic signature in the rhizome compared to the rest of the plant, a possible indication of nitrogen from an atmospheric source (Fig. S1), though *in situ* experiments with stable isotope tracers are needed to confirm the presence of nitrogen fixation.

Nitrogen fixation by microbial associates provides a key means of increasing the availability of ammonium, possibly supporting primary productivity. *P. scouleri* biomass reaches 12.7 kg of wet mass per square meter of shore and exudes 0.93 mg C per hour per gram dry mass as dissolved organic carbon that may fuel microbial activity (59). There is evidence that nitrogen fixation can contribute to seagrass productivity (66, 84), a possible adaptation to low nitrogen environments. Our finding that nitrogen fixing microbes are associated with a rocky intertidal surfgrass is especially surprising given that Tatoosh Island is in an area of upwelling and high DIN (86) at the more northerly end of the California Current Large Marine Ecosystem. Whether nitrogen fixation forms the basis for reciprocal host-microbe exchange is still unknown.

The metagenomic analyses we present here suggest that phototroph-associated microbiomes may be involved in carbon, nitrogen and vitamin metabolisms important to their hosts, likely generating commensal or mutualistic interactions. Future experiments should test these hypothesized interactions between host and microbiome. The importance of seaweeds and seagrasses to coastal productivity, and the demonstrated sensitivity of both host and microbes to increasing temperatures and pH (11, 62, 85), pathogens (61), and other anthropogenic stressors, underline the importance of further studying phototroph-microbiome interactions.

## Acknowledgements

Our gratitude to the Makah Tribal Nation for access to Tatoosh Island. We thank The University of Chicago’s Microbiome Center for pilot award funding, and Washington Department of Natural Resources grants 93099282, 93100399 (CAP) and NSF-DEB grant (#1556874) awarded to JT Wootton. We appreciate the work of C Sauceda in the isotope analysis, and A Wootton, A Wood and K Foreman in the field sampling. KM was supported by an EE Fellowship from The University of Chicago. S Owens and S Greenwald at Argonne National Lab provided expertise in sequencing.

## Supplementary Files

**Table S1**. A summary of eight metagenomes from five macrophyte taxa.

**Table S2**. A summary of taxonomy of MAGs

**Table S3**. The features of 72 metagenome assembled genomes (MAGs).

**Table S4**. Genes used to generate Fig. 3

**Table S5**. The features of *nifH* genes found in MAGs.

**Table S6**. *nifH* reference amino acid sequences

**Appendix 1.** Additional methods to quantify carbon and nitrogen stable isotopes in *P. scouleri*

**Figure S1**. Stable isotope analysis of delC13 and delN15 at blade tip, meristem, rhizome of P. scouleri. From blade tip to rhizome, water flow and thus elemental mixing reduces due to attenuation and boundary layer effects of surfgrass canopy. Assuming elemental uptake occurs from the same pools of C and N, the lower the extent of mixing, the heavier the isotopic signature should be at that point of the plant. This is observed with delC13 which gets heavier from the tip to the blade. This is observed with delN15 till the meristem after which it lightens. This is probably occurring as nitrogen is taken up from a different pool of nitrogen from that around the blade/meristem. This different pool is probably made available through n-fixation

